# A Fully Defined Synthetic Medium Mimicking Sugar Cane Molasses

**DOI:** 10.1101/2023.01.27.525923

**Authors:** Kevy Pontes Eliodório, Gabriel Caetano de Gois e Cunha, Felipe Senne de Oliveira Lino, Morten Otto Alexander Sommer, Andreas Karoly Gombert, Reinaldo Giudici, Thiago Olitta Basso

**Affiliations:** Department of Chemical Engineering, Escola Politécnica, Universidade de São Paulo, Av. Prof. Luciano Gualberto, 380, 05508-010, São Paulo, Brazil; Nosh.bio GmbH, Schwarzschildstraβe 6, 12489 Berlin, Germany; Novo Nordisk Foundation Center for Biosustainability, Technical University of Denmark, 2800, Kongens Lyngby, Denmark; School of Food Engineering, University of Campinas, R. Monteiro Lobato 80, 13083-862, Campinas, Brazil

**Author notes:** Corresponding author, Email address (Basso, T.O.), Postal address: Av. Professor Lineu Prestes 580, 05508-000, São Paulo - SP, Brazil. both authors contributed equally to this work (Kevy Pontes Eliodório, Gabriel Caetano de Gois e Cunha).

**Keywords:** Yeast fermentation, Defined medium, Synthetic molasses, Yeast physiology, Industrial strains, Alcoholic fermentation, Industrial biotechnology

## Abstract

**Background:** Yeast research in the context of food/beverage production and industrial biotechnology faces a dilemma: to use real industrial media or to use fully defined laboratory media? While the former option might lead to experiments closer to industrial conditions, the latter has the advantage of allowing for reproducibility and comparability of results among different laboratories, as well as being suitable for the investigation of how different individual components affect microbial or process performance. It is undoubtable that the development of a synthetic must a few decades ago led to important advances in wine yeast research.

**Results:** We developed a fully defined medium that mimics sugarcane molasses, a frequently used medium in different industrial processes where yeast is cultivated. The medium, named 2SMol, builds upon a previously published semi-defined formulation and is conveniently prepared from some stock solutions: C-source, organic N, inorganic N, organic acids, trace elements, vitamins, Mg+K, and Ca. We validated the 2SMol recipe in a scaled-down sugarcane biorefinery model, comparing the performance of different yeast strains in different real molasses-based media. We also showcase the flexibility of the medium by investigating the effect of nitrogen availability on the ethanol yield during fermentation.

**Conclusions:** Here we present in detail the development of a fully defined synthetic molasses medium, and we hope the 2SMol formulation will be valuable to researchers both in academia and industry to obtain new insights and developments in industrial yeast biotechnology.

## BACKGROUND

Molasses is a broad term used to describe concentrated sugar cane or sugar beet juice solutions after sucrose crystals removal [1]. Consequently, molasses is not the primary outcome but an industrial by-product of the raw sugar production process. Molasses is produced by water evaporation from clarified juice and sucrose crystal separation via centrifugation. Molasses can be reprocessed to increase the extraction of sugar crystals, resulting in a higher salt-to-sugar ratio and poorer quality of the final molasses for further industrial applications [2].

Molasses is an inexpensive renewable substrate with applications in many bioprocesses, such as the production of bioethanol, butanol, citric acid, and lactic acid, among many others [3]. During bioethanol production in Brazilian sugarcane-based biorefineries, molasses plays a unique role and significantly impacts production costs [4]. In this country, bioethanol production is traditionally coupled with sugar industries, thus enabling biofuel production from a must prepared from sugar cane juice and/or molasses [5]. The coupled production allows for flexibility and decreases the commercial risk associated with price fluctuation in sugar and ethanol. Still, it also affects the fermentation medium, or must [6].

Sugarcane molasses contains high amounts of fermentable sugars and other nutrients. Various compounds, including growth factors, macro and micronutrients, such as trace elements and vitamins, vary depending on the sugar cane variety, soil, climate, and processing conditions [4, 7, 8]. Additionally, molasses contains high concentrations of mineral compounds and salts, low nitrogen concentrations, and high concentrations of some compounds generated during sugar cane processing that might inhibit yeast performance [4, 5].

Molasses composition can affect both microbial growth and ethanol production. However, how the composition affects microbial physiology and fermentation performance is not yet completely clear, sometimes leading to low ethanol yields and high cell viability loss [3, 9]. This is partially due to the difficulties in achieving reproducibility in such studies because molasses composition is so variable, and its composition is rarely determined before physiological studies at industrially relevant conditions are carried out [3, 7, 10]. Furthermore, evaluating microbial performance in such media is of pivotal importance in selecting improved yeast strains for industrial fermentations [11].

A solution to increase reproducibility among experiments carried out with different molasses and to obtain a better understanding of microbial performance under industrially-relevant conditions is to develop a fully defined synthetic medium that summarizes the main properties of real molasses. Commonly used laboratory media for physiological studies (e.g., YNB, YPD, Mineral media, among others) are not able to properly mimic industrial conditions, especially in the case of sugar cane molasses [3]. For this reason, previous studies attempted to formulate synthetic media, chemically defined or complex, to simulate many industrial substrates, such as lignocellulosic hydrolysates, sugar cane molasses, malt wort, and grape juice, which is probably the most relevant in the context of such synthetic media [3, 12 – 16].

In fact, wine fermentation research has used synthetic grape juice since the 1990s with different formulations based on specific features of each grape. Bely et al. [12] proposed a synthetic medium to simulate a standard grape juice while evaluating the effects of assimilable nitrogen in the kinetics of yeast fermentations. Establishing a basal synthetic medium mimics a natural medium used for wine fermentation and facilitates investigating the effect of compositional changes in the process.

Regarding sugarcane molasses media, Chandrasena et al. [13] proposed a *“synthetic molasses medium”* during their investigation on the effects of metal ions in beer fermentation. However, the reported results for the proposed synthetic wort did not reproduce the ethanol yield observed with the industrial substrate. Lino et al. [3] recently developed a semi-synthetic sugar cane molasses medium. The medium was developed by adjusting the Carbon/Nitrogen ratio and Phosphorus, Potassium, Magnesium, and Calcium levels. In addition, other compounds were added, such as malic acid and trans-aconitic acid (organic acids present in sugarcane juice). The Maillard reaction products were also included by simulating the reaction between sugars and amino acids at high temperatures during the preparation procedure. The selected nitrogen sources were amino acids (mostly consumed during the Maillard reaction), ammonium salts, and peptone, which led to a final complex formulation. The reported semisynthetic medium mimicked the results of industrial sugar cane molasses media accurately under fed-batch operation in bench-scale ethanol production. To our knowledge, it is the closest result of a Brazilian-like sugarcane molasses fermentation reported. Although the authors have found good agreement between this semi-defined synthetic medium and industrial molasses based-media, the presence of peptone and the use of Maillard reactions for its formulation, could hamper reproducibility among different laboratories. In addition to that, concentrations of some relevant ions in synthetic molasses medium (such as K^+^ and Ca^2+^) are different from reported data on industrial molasses in fuel ethanol fermentation [4].

Therefore, here we aimed to conduct an extensive investigation based on the reported semi-synthetic molasses proposed by Lino et al. [3], studying the effects of the main components on yeast physiology. With these results, we propose here a fully defined synthetic molasses medium, without addition of complex ingredients and with the flexibility to be prepared with different nutrient concentrations. The proposed defined medium was benchmarked against three different Brazilian sugarcane molasses in industrial-like fermentations. Finally, a case study is presented to exemplify how this medium can aid in performing scientific research.

## METHODS

### Industrial molasses samples and chemical composition analysis

Industrial molasses samples were gently provided by five sugar cane biorefineries from São Paulo State (Brazil) in different collection dates ranging from 2014 to 2019. The molasses samples were kept in 5 °C refrigerators and, before use, were diluted with distilled water to yield approximately 500 g of molasses.L^-1^. The resulting diluted molasses were centrifuged (2000 g, 15 min) to remove insoluble materials which affect analytical methods [11]. Next, the refractive index of the supernatant was adjusted to 20° Brix with a handheld refractometer. Molasses was then heat-sterilized at 121 °C for 20 min. Before all experiments, the total reducing sugar (TRS) concentration was measured using high-performance liquid chromatography (HPLC).

In order to compare the results for the proposed defined synthetic molasses with real sugarcane molasses samples, the main nutrients (minerals and nitrogen, phosphorus and sulphur sources) of three industrial molasses (Mol_A, Mol_B, and Mol_C) were determined in an outsourced company (ALS Ambiental Ltda., São Paulo, Brazil). The chemical compounds and their respective methods were the following: Total nitrogen (SMWW 23a Ed. 2017 - method 4500 N C); Sulphur (SMWW 22a Ed. 2012 - 3120 B); Potassium, Magnesium, Calcium, Phosphorus, Sodium, Zinc, Manganese, and Copper (USEPA 6020 A); Chloride (USEPA 9056 A: 2007, 300.1: 1997).

### S. cerevisiae strains used and their preservation

Industrial bioethanol strain PE-2 [5] and the laboratory haploid strain CEN.PK113-7D were gently provided by Dr. Luiz Carlos Basso (*University of São Paulo, Escola Superior de Agricultura Luiz de Queiroz, Piracicaba, SP, Brazil*). Each strain was grown overnight (30 °C, 200 rpm) in YPD medium (1% yeast extract, 2% bacteriological peptone, and 2% glucose). After growth, glycerol was added to achieve a final concentration of 20% v/v. Cell aliquots were then stocked in cryotubes at – 80°C [17].

### Semi-synthetic molasses preparation

Preparation of the semi-synthetic molasses (named LSM) was performed according to Lino et al. [3] with some modifications. Three stock solutions were prepared with phosphate buffer saline (PBS, pH 7.4): a solution with sugars and amino acids, a solution with nutrients with concentrations higher than 0.1 g.L^-1^, and a final solution with nutrients with concentrations lower than 0.1 g.L^-1^. Here, the macronutrients solution and the micronutrients solution were divided into two solutions each.

The proposed 5-fold concentrated macronutrient solution was split into a 5-fold concentrated broth stock (in g.L^-1^): peptone, 24.5; (NH_4_)_2_SO_4_, 0.5; (NH_4_)_2_PO_4_.4H_2_O, 7.1; NaCl, 2.5; MgSO_4_.7H_2_O, 5.01; CaCl_2_.2H_2_O, 0.336; and KCl, 0.06; and a 5-fold concentrated organic acids solution (in g.L^-1^): trans-aconitic acid, 10.0; L-malic acid, 5; and citric acid, 0.05. Both stock solutions were heat-sterilized at 121 °C for 15 min. The micronutrients and growth factors were split into a vitamin stock solution 100-fold concentrated containing a final concentration in the semi-synthetic molasses (mg.L^-1^): inositol, 10; nicotinic acid, 10; calcium pantothenate, 1; biotin, 0.01; pyridoxine hydrochloride, 0.04; thiamine hydrochloride, 0.04; and para-aminobenzoic acid, 2; and trace elements stock solution 100-fold concentrated (mg.L^-1^):0.5 H_3_BO_3_, 0.5;4 MnSO_4_.H_2_O, 0.4; ZnSO_4_.7H_2_O, 0.4; FeCl_3_.6H_2_O, 17; Na_2_MoO_4_.H_2_O, 31; KI, 12, and CuSO_4_.5H_2_O, 0.4. Both solutions were filtered sterilized (0.22 μm).

### Preparation of the fully defined synthetic molasses

Preparation of the medium developed in this work (named 2SMol) was based on mixing seven sterilized stock solutions. Initially, the sugar stock solution described by Lino et al. [3] was mixed in a 0.25 ratio to a filter-sterilized (0.22 μm) 500 g.L^-1^ sucrose solution to yield a new sugar stock solution containing 80% of sucrose. This procedure was adopted because a previous investigation indicated partial hydrolysis of sucrose in LSM caused by heat sterilization. Later, the procedure was further adapted by mixing the sugar stock with a filter-sterilized amino acid solution. For this purpose, a solution containing sucrose (400 g.L^-1^), glucose (50 g.L^-1^), and fructose (50 g.L^-1^) was filter-sterilized, generating the new sugar stock solution. Next, the amino acid solution was prepared by adding aspartic acid (3.75 g.L^-1^), glutamine (10.375 g.L^-1^), and asparagine (3.55 g.L^-1^), with pH adjustment using H_2_SO_4_ (1 M) to 5.0, followed by a filtering process.

Vitamins and trace elements solutions were prepared according to Verduyn et al. [18]. The salt stock solution (K_2_SO_4_, MgSO_4_), the organic acids stock solution (trans-aconitic, L-malic, and citric acids, and KOH), and the Calcium stock solution (CaCl_2_) were heat sterilized at 121 °C for 15 min. The inorganic nitrogen ((NH_4_)2HPO_4_) stock solution was filter-sterilized (0.22 μm) to avoid thermal degradation.

The final composition of the medium was adjusted along the experiments by changing the proportions of the stock solutions mentioned above, as described in the Results section when appropriate.

### Yeast growth kinetics in microtiter plates

Growth profiles and parameters were used to compare real molasses and the synthetic medium developed (2SMol). Microplate assays were employed in this step. Inocula for the microplate assays were prepared by inoculating a loopful of yeast cells from a cryotube culture into YPD medium (1% yeast extract, 2% peptone, and 2% glucose) overnight (30°C, 200 rpm). On the next day, 0.1 mL of the cell suspension was transferred to a fresh YPD medium, incubated in the same conditions above, and monitored by the optical density at 600 nm (OD_600_) [19]. Cells were harvested during the exponential phase and carefully washed five times with sterile distilled water to minimize micronutrients interference from the YPD medium. Finally, 20 μL of the washed cell suspension were incubated in 180 μL of each media (initial OD_600nm_ of 1) in a microplate reader Infinite M200PRO (Tecan, Switzerland) and growth was monitored by measuring OD_600_ in 20 min intervals at 30 °C. All conditions were performed in triplicates. Blank was prepared by incubating 180 μL of each media with 20 uL of pure sterile water.

Besides visual inspection, three parameters were used to numerically compare the growth profiles: the maximum specific growth rate (μ), the maximum OD_600nm_ (OD_max_), and the deceleration time (t_decel_). μ is determined as the highest slope in the ln OD_600nm_ versus time curve. OD_max_ is the maximum OD_600nm_ reached, representing total cell growth. The additional parameter t_decel_ was defined as the time interval between the end of the exponential growth phase and the time to reach OD_max_. A script was developed in Python to determine all three parameters in an automatic manner.

### Substrate and extracellular metabolites quantification

A HPLC system (Shimadzu Prominence LC-20AB, Japan) with a refractive index detector was used to determine sugars and metabolites concentrations in fermentation experiments. Before injection, liquid samples were centrifuged (10,000 rpm, 10 minutes) to remove suspended solids or cells. The supernatant was 10-fold diluted to match the HPLC limits of quantification and filtered (0.22 μm syringe filter). Linear calibration for each component was performed for each injection batch.

Two different columns were used. A Bio-Rad HPX-87H column at 60°C was used to separate glucose, fructose, glycerol, and ethanol in fermentation samples with 5 mM H_2_SO_4_ as eluant (0.6 mL.min^-1^). Sugar concentrations (sucrose, glucose, and fructose) in molasses and medium samples were determined using a Bio-Rad HPX-87C column kept at 85 °C with ultrapure water (0.6 mL.min^-1^) as the eluent. The injection volume was 10 μL for both columns.

### Statistical analysis

Minitab 19 (Minitab Inc., USA, version 19.2–64 bit) was used to perform all statistical analyses. During the individual investigation of each nutritional group, multiple comparison tests for each evaluated parameter were performed by one-way ANOVA and Tukey’s and Games-Howell’s tests for homo and heteroscedastic data, respectively [20, 21]. Bartlett statistical test (p<0.05) was applied to the variances test [21]. After this initial investigation, the statistical software was used to define a 3^3^ complete factorial design, yielding 27 conditions, to evaluate the interactions among the concentration of amino acids, ammonium, and vitamins on μ, OD_max_, and t_decel_ for the *S. cerevisiae* PE-2 and CEN.PK113-7D strains. This methodology allowed for the creation and analysis of a statistical model given in Equation 1 for the coded variables also performed in Minitab. The response, linear, and interaction terms are represented by y, bi, and bij, respectively:

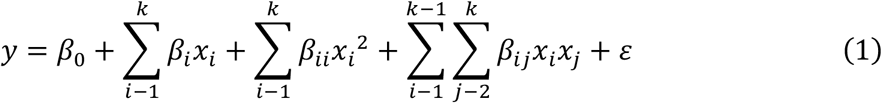

### Validation of the synthetic molasses and case study in a scaled-down sugar cane biorefinery

After microplate experiments, the final composition of 2SMol was benchmarked against three real molasses samples from different mills (São Paulo, Brazil) named Mol_A, Mol_B, and Mol_C in experiments mimicking Brazilian ethanol production. The protocol was described in detail by Raghavendran et al. [11]. This protocol accounts for many specific features of Brazilian ethanol production, such as cell recycling, sequential fermentation, acid treatment, and non-aseptic conditions. The industrial strain of *S. cerevisiae* PE-2 and 32°C were selected for this experiment. The same protocol and conditions were applied to the case study investigating the total nitrogen concentration for the media 0SMol, 2SMol, and 4SMol.

## RESULTS AND DISCUSSION

### A fully defined synthetic medium to mimic industrial sugarcane molasses

Reproducing the relevant conditions of industrial substrates in the laboratory can be challenging due to the wide compositional variation and eventual lack of information regarding some specific compounds in such substrates [43]. Accordingly, reproducibility among different laboratories is often compromised [3]. In the sugarcane fermentation industry, a synthetic medium to mimic this substrate should consider a basal composition with nutrients required for yeasts. It should also feature special conditions such as high salt concentrations and the presence of inhibitory compounds.

In order to obtain a fully defined chemical composition reproducing sugarcane molasses, we took the composition reported by Lino et al. [3] as a starting point and adjusted it using data on real molasses compositions described in the literature [2, 4]. The defined composition of our medium, which we named 2SMol, is presented in Table 1.

**Table 1.**
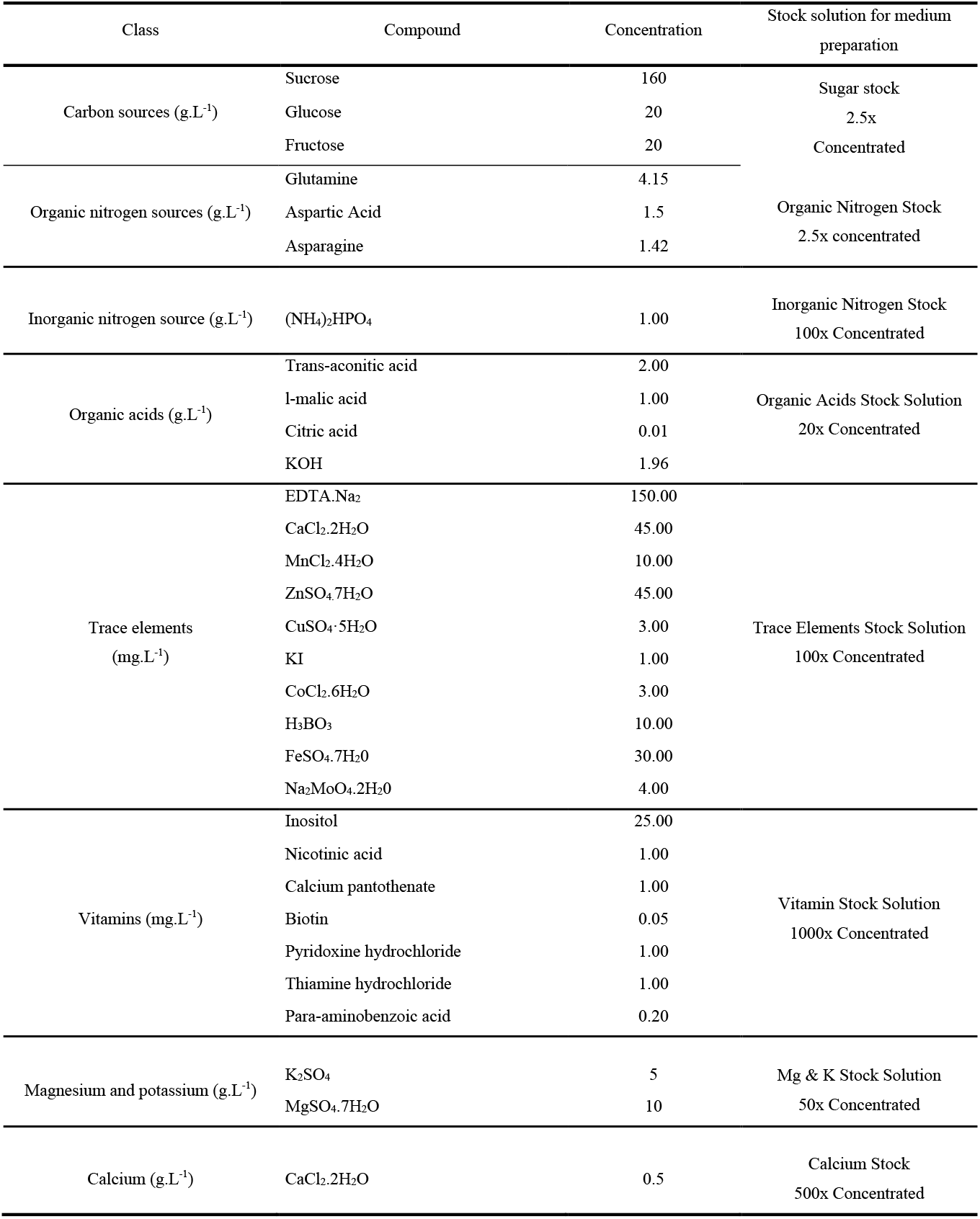
Composition of the fully defined synthetic molasses medium (2SMol).

The major modifications to the composition reported by Lino et al. [3], used to formulate 2SMol, and the respective rationales are the following:

- The sugar proportion was adjusted with a filter-sterilized 2.5-fold concentrated solution. The proportion was set to sucrose at 80%, fructose plus glucose at 10% each (in w/v) to match reported values from real-world molasses samples (Figure S1, Supplementary data 1);
- Vitamin and trace elements stock solutions were based on the widely used defined medium reported by Verduyn et al. [18], commonly adopted for studies in quantitative yeast physiology. Vitamin concentrations are set according to the original medium [18], and trace elements were 10-fold increased to approximate literature data for sugarcane molasses [2, 4];
- Peptone addition, as reported by Lino et al. [3], was removed, since it represents a complex source of nitrogen and of other nutrients, the composition of which if not fully defined. Instead, inorganic nitrogen ((NH_4_)_2_HPO_4_) and organic nitrogen (amino acids) sources, provided as filter-sterilized solutions (to avoid thermal degradation), were included;
- The organic acids solution proposed by Lino et al. [3] was kept here, but with the addition of citric acid [7] and pH adjustment using KOH, leading to a final pH of 5.0; the potassium ions added for pH adjustment were accounted for in the final composition;
- Calcium (as calcium chloride), magnesium and potassium ions (as sulphate salts) were split into two different stock solutions since they can differently affect yeast metabolism [22].

The first synthetic molasses formulation (named 1SMol) was based on the combined sugar and organic nitrogen source stock solution described by Lino et al. [3] (Methods). Growth kinetics experiments in microplates showed that the growth profiles of yeast in this 1SMol medium were very different from growth profiles obtained using real industrial molasses media, even when supplemented with increasing amounts of vitamins and inorganic nitrogen sources, two of the leading nutritional groups affecting growth (data not shown). In addition, as mentioned before, the organic nitrogen sources were combined with sugars in the 1SMol formulation, meaning that there was no flexibility in adjusting the nitrogen concentration independently from the TRS concentration. As the preparation of this stock solution involved heat-sterilizing of amino acids and sugars, caramelization and/or Maillard reactions occurred [23], leading to the degradation of amino acids and inversion of sucrose into glucose and fructose. Indeed, the investigation of the sugar composition of the synthetic molasses proposed by Lino et al. [3] indicated a sucrose proportion of only 10%, which is much lower than the values observed for true molasses samples (81 ± 6%) (Figure S1, Supplementary data). Moreover, these degradation reactions can result in unknown compounds in the medium, resulting in a non-defined medium composition.

To avoid these issues, the 2SMol formulation included an individual filter-sterilized organic nitrogen solution containing amino acids in concentrations commonly found in sugarcane substrates [24]. This modification avoided the formation of unknown reaction compounds and enabled the investigation of the influence of individual amino acids on yeast performance. Although yeasts can grow without organic nitrogen sources, amino acids can save sugar carbon and energy in yeast’s metabolism, as evidenced by Albers et al. [25].

Diammonium hydrogen phosphate [(NH_4_)_2_HPO_4_] was added to the medium as an inorganic nitrogen source, playing a fundamental role in the biosynthesis of amino acids for biomass formation in *S. cerevisiae* [25]. In addition, this solution is also the primary source of phosphorous, required for nucleic acid synthesis, and as a substrate for many enzymes [26].

The most common organic acids found in sugarcane were added in accordance with the report from Lino et al. [3]. Weak organic acids are known to cause an inhibition effect on yeast cells since they permeate through the cell membrane in the protonated form, dissociating in the yeast’s cytosol. As a result, cells excrete one H^+^ using 1 ATP, to maintain intracellular pH homeostasis, decreasing the final biomass yield [4, 27]. Organic acids are also known to cause decreased DNA and RNA synthesis rates and diminished metabolic activity [28].

Salts were divided into two solutions and added in the range of concentrations reported by Basso et al. [4]. According to these authors, K, Mg, and S are the most abundant elements in molasses, apart from for C, H, and O. Thus, the concentration of these elements was increased when compared to those employed by Lino et al. [3]. Figure 1 illustrates the concentration ranges of the main chemical elements, as reported by Basso et al. [4] (lower and upper limits), in comparison with the 2SMol formulation, with the values reported by Lino et al. [3] and with values measured in real molasses (Mol_A, Mol_B, and Mol_C) from sugarcane biorefineries, all located in the state of São Paulo, Brazil.

**Figure 1.**
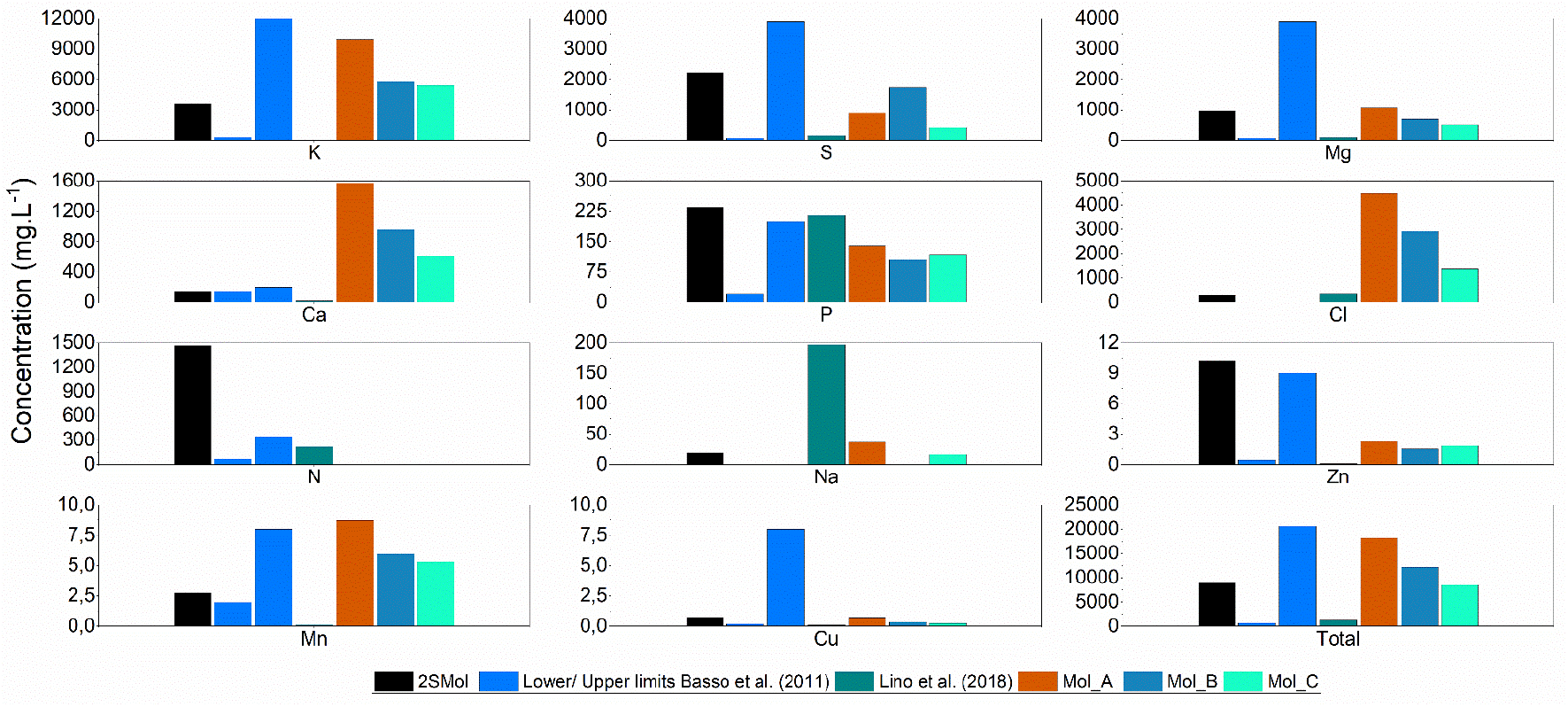
Concentrations of different chemical elements in the 2SMol formulation, compared to data reported by Basso et al. [4], Lino et al. [3], and to real sugarcane molasses media obtained from biorefineries Mol_A, Mol_B, and Mol_C.

Calcium was added to achieve the lower limit reported [2, 4], due to its detrimental effect on yeast since it acts as a flocculation facilitator and as a suppressor of magnesium-dependent enzymes [22]. Other mineral compounds, such as Zn, Mn, and Cu, were adjusted to the range values by a 10-fold increase in the trace elements solution proposed by Verduyn et al. [18]. According to Figure 1, zinc (from trace elements solution), phosphorous (inorganic nitrogen source) and nitrogen are close to the upper reported limit. A proper evaluation of each nutritional group may clarify their respective effects on growth.

### Adjustment of amino acids concentrations

Within all compounds in 2SMol, the nitrogen source (as a sum of inorganic and organic sources) presented values up to 4 times higher (1.47 g.L^-1^) than the values reported previously (0.35 g.L^-1^) (Figure 1). As already discussed, this nutritional source presents an important effect on *S. cerevisiae*’s physiology. Therefore, we proposed an additional experiment in microplates to evaluate yeast growth (using two strains, *S. cerevisiae* PE-2 and CEN.PK113-7D) at different organic nitrogen concentrations. The new versions of the synthetic medium were developed by altering the amino acid concentration with respect to the 2SMol-basal medium (0, 25, 50, 75, 100% of the 2SMol original organic nitrogen composition). The results for the growth parameters μ, OD_max_, and t_decel_ were benchmarked against industrial molasses media (Mol_A, Mol_B, and Mol_D). The results for parameters and statistical groups are presented in Table 2.

**Table 2.** Maximum specific growth rate, maximum optical density, and deceleration time for the growth of *S. cerevisiae* on 2SMol compositions with different amino acid concentrations or on industrial molasses.

| Condition | $\mu$ (h<sup>-1</sup>) | | ODmax | | tdecel (h) | |
| --- | --- | --- | --- | --- | --- | --- |
|  | PE-2* | CEN.PK113-7D† | PE-2† | CEN.PK113-7D† | PE-2* | CEN.PK113-7D* |
| AA 100% | 0,420 ± 0,001 | 0,363 ± 0,004 | 0,919 ± 0,012 | 0,912 ± 0,010 | 8,67 ± 0,00 | 11,67 ± 0,58 |
|  | B | AB | A | A | D | D |
| AA 75% | 0,429 ± 0,00 | 0,360 ± 0,00 | 0,910 ± 0,018 | 0,900 ± 0,014 | 8,89 ± 0,19 | 14,34 ± 0,33 |
|  | B | B | A | A | D | D |
| AA 50% | 0,427 ± 0,00 | 0,360 ± 0,00 | 0,859 ± 0,019 | 0,877 ± 0,021 | 8,67 ± 0,00 | 16,34 ± 2,60 |
|  | B | AB | AB | A | D | BCD |
| AA 25% | 0,404 ± 0,01 | 0,360 ± 0,01 | 0,813 ± 0,018 | 0,801 ± 0,017 | 11,78 ± 0,19 | 19,78 ± 0,39 |
|  | B | B | BC | B | C | C |
| AA 0% | 0,349 ± 0,00 | 0,310 ± 0,00 | 0,682 ± 0,017 | 0,667 ± 0,007 | 24,89 ± 1,50 | 33,12 ± 1,26 |
|  | C | C | D | D | A | A |
| Mol_B | 0,496 ± 0,01 | 0,390 ± 0,02 | 0,902 ± 0,015 | 0,827 ± 0,016 | 5,78 ± 0,19 | 10,78 ± 0,19 |
|  | A | A | A | B | E | D |
| Mol_A | 0,426 ± 0,011 | 0,370 ± 0,01 | 0,682 ± 0,057 | 0,666 ± 0,019 | 8,56 ± 0,19 | 14,56 ± 0,51 |
|  | B | AB | D | D | D | D |
| Mol_D | 0,417 ± 0,010 | 0,380 ± 0,01 | 0,746 ± 0,009 | 0,716 ± 0,017 | 17,56 ± 0,77 | 26,00 ± 0,88 |
|  | B | AB | CD | C | B | B |

\*Pairwise comparisons using Games-Howell test $\alpha=0,05$ .
†Pairwise comparisons using Tukey test with $\alpha=0,05$ .

In general, higher values of μ were obtained for PE-2 when compared to CEN.PK 113-7D strain. Among all amino acids concentrations tested, statistically different results were only observed for the absence of organic nitrogen (0% AA, group C). Among industrial samples, Mol_B was statistically different from Mol_A and Mol_D. Additionally, the values for this parameter for Mol_A and Mol_D were close to the synthetic compositions, except for 0% of amino acids.

The results for OD_max_ were similar between the tested strains, with increasing values as amino acid concentration increased. In higher nitrogen concentrations (75 and 100% AA), similar results (0.92 ± 0.01 and 0.91 ± 0.02 for PE-2 and 0.91 ± 0.01 and 0.90 ± 0.01 for CEN.PK113-7D) were observed, indicating that the strains might be insensitive to the amino acid concentration in this range. The industrial molasses presented statistically different OD_max_ values for higher (75 and 100 %) and lower (0%) amino acids concentrations.

The decrease in amino acid concentrations led to increased deceleration times ranging from 8.7 to 24.9 and 11.7 to 33.1 h for PE-2 and CEN.PK113-7D, respectively. Deceleration times for PE-2 were significantly lower than CEN.PK113-7D for all tested media, although the pairwise comparison was not performed. For this parameter, the industrial molasses presented values comparable with the ones obtained for the synthetic compositions, except for Mol_B, which presented a statistically different value.

The growth kinetics obtained from this set of experiments indicated that at both 25 and 50% amino acid concentrations, the growth profiles fell within the range of the growth profiles observed for growth on molasses Mol_B and Mol_D, which represent the upper and lower limits, respectively, among the industrial samples investigated (Figure S2, Supplementary data). In the lower concentration (25%), the carbon-nitrogen ratio, considering carbon in fermentable sugars and total nitrogen, was 159, which is in the midrange of values reported in the literature for sugarcane media – from 57 to 209 [3, 29]. Therefore, the 25% amino acid concentration of 2SMol was selected for subsequent experiments, which aimed at testing the individual effects of all other nutritional groups.

### Assessing the effect of nutritional groups: inorganic and organic nitrogen and vitamin sources affect S. cerevisiae growth patterns

Variations to the 2SMol composition (Table S1, Supplementary data) were tested to assess the effects of each nutritional group on growth profiles and physiological parameters of *S. cerevisiae* strains PE-2 and CEN.PK113-7D. The effects of phosphate and inorganic nitrogen were separately studied by using ammonium sulphate and potassium phosphate (instead of diammonium hydrogen phosphate), keeping the levels of the complementary chemical elements (sulphur and potassium) the same as in the 2SMol original formula. It means, for example, that in the composition of zero ammonium, potassium phosphate was added to keep the level of phosphorus equal to the one found in the original formula.

Figure 2 depicts the growth curves obtained for the three main nutritional groups. The amino acids and ammonium concentrations were the most significant variables influencing the physiological parameters in both strains (Figure S3, Supplementary data). The effect of nitrogen sources (inorganic and organic) was statistically significant for all parameters for both strains (Table S2, Supplementary data), except for the parameter t_decel_ for *S.cerevisiae* CEN.PK113-7D cultivations. In fact, these two groups were expected to affect the evaluated parameters since yeast displays complex regulatory systems to adapt to nitrogen availability, strongly impacting alcoholic fermentation and growth kinetics [30].

**Figure 2.**
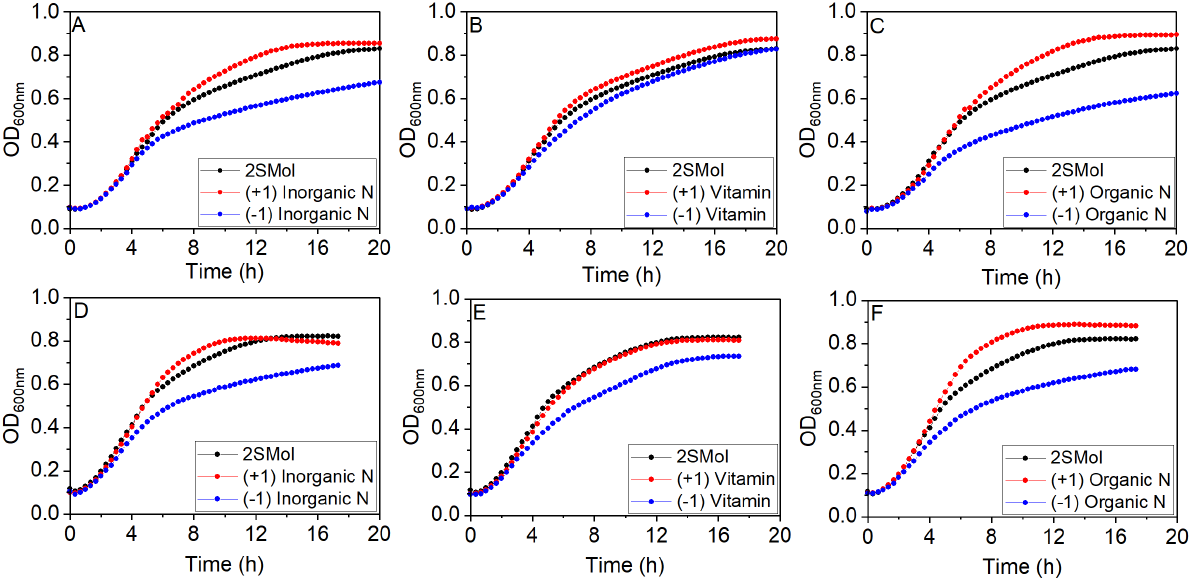
Growth curves for different versions of the defined synthetic molasses (2SMol) for two *S. cerevisiae* strains: CEN.PK113-7D (A, B, and C) and PE-2. (D, E, and F). (A) and (D) presents higher (red dots) and lower levels (blue dots) of inorganic nitrogen concentrations compared to 2SMol basal formulation (black dots); (B) and (E) presents higher and lower levels of vitamin concentrations, and (C) and (F) presents higher and lower levels of organic nitrogen concentration

Vitamins also had a significant effect on the growth of both the *S. cerevisiae* CEN.PK113-7D and PE-2 strains, in terms of the parameters μ and OD_max_, respectively. The vitamin concentration only had a statistically significant effect (p<0.05) on μ of the CEN.PK113-7D strain. On the other hand, the vitamin effect was smaller for the PE-2 strain, evidencing the capability of the PE-2 strain to adapt to low vitamin concentrations. Unexpectedly, vitamin concentration (−1) affected only PE-2 OD_max_ results, although the same final OD_max_ was evidenced for PE-2 and CEN.PK113-7D (Figure 2 B and E).

Besides OD_max_, some identified genetic signatures could explain the ability to cope with a decreased vitamin availability in industrial strains. Stambuk et al. [31] demonstrated that SA-1, PE-2, and other industrial fuel ethanol strains contain amplification of the *SNO2/SNZ2* and *SNO3/SNZ3* genes, two pairs involved in the biosynthesis of thiamine and pyridoxine (vitamins B1 and B6, respectively), which are present in the vitamin stock solutions used. Similarly, Argueso et al. [32] also detected five extra copies of the *SNO/SNZ* genes in a *S. cerevisiae* strain JAY270 (PE-2 derived) in comparison with the *S. cerevisiae* laboratory strain S288C. The expression level of these genes was up-regulated approximately 4-fold in the industrial strain relatively to the laboratory strain. The results presented by Stambuk et al. [31] and Argueso et al. [32] were recently clarified by Paxhias and Downs [33]. It was demonstrated that *SNZ2* and *SNZ3* genes were sufficient to generate growth on a minimal vitamin medium (only biotin and D-pantothenic acid hemicalcium salt were added as vitamins).

Organic nitrogen, inorganic nitrogen and vitamins were chosen for a subsequent study. For this purpose, a factorial design with three levels (Table S3, Supplementary data) was used to evaluate the interactions among these variables and establish a statistical model to emulate different molasses with these parameters. Additionally, two industrial molasses samples were tested in the same experimental runs, comparing their results to the model obtained from cultivations using the 2SMol medium. Each condition was run in triplicate. The investigated ranges were the same tested for inorganic nitrogen in the previous experiment, with organic nitrogen concentration increased to cover a vaster dimensional space.

Application of the ANOVA (Table S4-6, Supplementary data) for the three variables indicated that the statistical model described all growth parameters very well. The high R^2^ values for the adjusted quadratic model displayed an overall good fit. The R^2^ values were close to 90% for almost all parameters in both strains, indicating a suitable model. Additionally, high t-values and low p-values also supported the statistical significance of the regression coefficients [20].

The investigation of vitamins, inorganic and organic nitrogen resulted in a more robust description of the effects of these essential nutritional groups for *S. cerevisiae* growth, not obtained in the first investigation of this section. The obtained equations could then be used, for example, to estimate the concentration ranges that would theoretically reproduce the parameters of distinct molasses samples.

Finally, the growth parameters obtained for the synthetic compositions tested agreed with those found in industrial molasses (Figure S4, Supplementary data), indicating that the synthetic medium presents flexibility and sensitivity. In spite of the large variability in molasses composition, the results of this section indicate that the formulation of 2SMol might be adequate to represent real industrial samples and could be an important tool for alcoholic fermentation research.

### Validation assays performed in a scaled-down sugarcane biorefinery

Although microplate assays are a valuable tool for investigating many conditions and for defining the 2SMol composition, the proposed synthetic molasses was further validated in experiments more representative of industrial ethanol production.

For this purpose, the 2SMol formulation (37.5% of the amino acids) was used as substrate in this simplified scaled-down process. The results over five fermentation cycles were compared to those obtained with real industrial molasses samples (Mol_A, Mol_B, and Mol_C). All fermentation media were diluted to contain 180 g.L^-1^ TRS, and the experiments were performed in triplicates.

Initially, the total mass of each tube along the fermentation assay was registered hourly. The mass difference is associated with CO_2_ loss, directly correlated with the sugar consumption and ethanol production profiles. The profiles for 2SMol (Figure 3A) were consistent with those observed using industrial molasses, showing similar results. The results are presented as mmol CO_2.gwet biomass_^-1^ to account for possible differences associated with biomass variations along the cycles.

**Figure 3.**
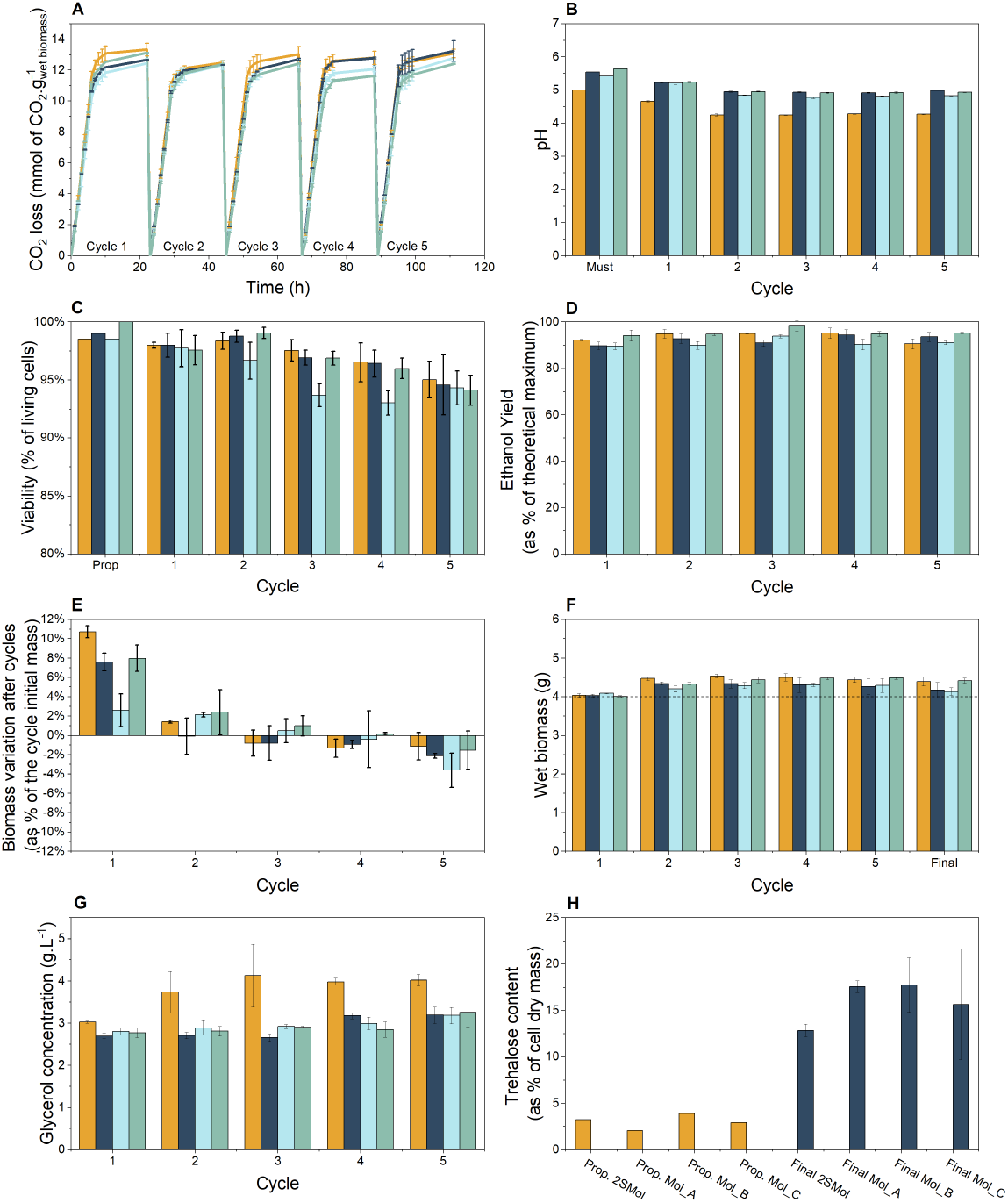
Performance of *S. cerevisiae* PE-2 in the defined synthetic molasses (2SMol) and in real sugarcane molasses media in a scaled-down sugarcane biorefinery throughout five consecutive fermentation cycles. The fermentations were performed in triplicates. The initial yeast biomass was 4 g (wet mass), acid treatment (pH set to 2.5, 1 h incubation at room temperature) was performed before each new cycle of fermentation. The tubes were incubated at 32 °C for 10 h and weighted hourly. The tubes were kept at room temperature (c.a. 20°C) overnight until the beginning of the next cycle. Orange (2SMol); Dark blue (Mol_A); Light blue (Mol_B); Green (Mol_C). (A) CO_2_ profiles (CO_2_ loss in mmol.g of wet biomass^-1^) over five fermentation cycles. (B) pH profiles of the media and fermentation wines. (C) Viability (as percentage of living cells, %) values by the methylene blue staining technique [44]. (D) Ethanol yield (as percentage of theoretical maximum 0.511 g of ethanol.g of TRS^-1^, %) calculated based on [11] and [35]. (E) Biomass variation (%) represented as percentage of the cycle initial wet yeast mass. (F) Overall biomass accumulation (g of wet biomass). (G) Glycerol concentrations (g.L^-1^). (H) Trehalose content (as % of cell dry mass) after propagation (orange) and fifth fermentation cycle (marine blue).

Although all media presented the same CO_2_ production rate (the slope of the curve), 2SMol presented higher CO_2_ loss at the end of first cycle. Indistinguishable results were observed for the second and third cycles. Differences were observed in the following cycles, in which the CO_2_ profile in the 2SMol resembled the profiles obtained using molasses from Mol_A, which reached higher values at the end of the cycle than Mol_B and Mol_C. The total CO_2_ loss at the end of fermentations remained between 12-14 mmol.g_wetbiomass_^-1^ in molasses (TRS concentration 19.4%) at 30 °C, agreeing with reported values for PE-2 [11].

The pH is vital to ensure fermentation efficiency and maintain cell viability. *S. cerevisiae* is an acidophilic organism with an optimal pH range for growth between 4 and 6 [34]. Thus, this parameter was measured in the fermentation media and wines (Figure 3B). Comparing fermentation media, 2SMol presented a final pH of 5, lower than 5.5 obtained for industrial molasses. For the fermentation cycles, a similar wine-pH pattern was observed with a decrease in the first (6.2 ± 1.6 %) and the second cycles (6.7 ± 1.7 %) followed by a stabilization around a constant value for all tested media.

These results were compared to the yeast viability profiles to assess if the differences in absolute values for pH were critical for the process. Figure 3C shows the viabilities measured by the methylene-blue method along the fermentation cycles. The obtained results were similar, with relatively high viability after propagation followed by a slight decrease in the cycles. Additionally, viability remained higher than 90% for all media, indicating that the pH differences for 2SMol were within a safe range for ethanolic fermentation. Moreover, if one may be interested, 2SMol can be easily adjusted by adding KOH during the formulation to match an exact demanded pH of a specific sample.

Yeast biomass is an essential parameter, reflecting nutritional conditions imposed on cells during fermentation. Literature suggests that the nitrogen content in sugarcane-based media is only sufficient to support an increase in biomass around 5 to 10% in relation to the beginning of each cycle [9]. This increase can replace cell loss during the process (e.g., during centrifugation or acid treatment to cell recycling), but it is routinely necessary to withdraw excess cell biomass from time to time in industrial scale [9].

In order to investigate if 2SMol could present similar results to industrial molasses, the biomass variation along the cycles was measured (Figure 3F). The results indicate that the synthetic medium presented similar behavior to the industrial molasses, increasing biomass in the first two cycles and no increase or decrease in the final one (Figure 3E). Interestingly, the biomass variation reported by Raghavendran et al. [11] indicated that PE-2 could increase biomass in all six cycles tested, different than the situation observed in our experiment. These small differences could be due to the difficulties in precisely measuring wet biomass.

Ethanol yield may be regarded as the most critical parameter in industrial fermentation, which directly relates to the economic viability of the process [35]. This parameter is represented here as a percentage of the maximum theoretical (or stoichiometric) yield, based on TRS fed, and details on its calculation are described elsewhere [11, 35]. The synthetic molasses composition (2SMol) displayed similar values to the ones obtained with real industrial molasses (Figure 3D). The values presented fluctuate and reach figures as high as 94% and no particular trend could be observed for any single type of medium used.

Another important metabolite we measured during our fermentation assays was glycerol, which represents the main by-product of ethanolic fermentation, in terms of carbon utilization. Its intracellular accumulation acts as a vital osmolyte, attributing protective properties against hyperosmotic and thermal stresses [36 – 38]. Among all measurements made here, glycerol production was the only output that resulted in significant differences when the 2SMol medium was compared to real molasses samples (Figure 3G).

While glycerol in wine from industrial molasses remained at approximately 3 g.L^-1^, the use of 2SMol led to concentrations around 4 g.L^-1^. Raghavendran et al. [11] report a glycerol concentration of approximately 4 g.L^-1^ for PE-2 in molasses at 30 °C. Additionally, Prado et al. [39] obtained glycerol titters higher than 4.5 g.L^-1^ for a thermo-tolerant strain grown at 34°C. Further investigations adjusting the composition of our 2SMol medium (e.g., lower salt concentrations) or in different osmotic pressures can potentially result in similar glycerol concentrations, with similar biomass variation, ethanol yields, and viability.

Trehalose also protects cells against stressful environments [40]. Thus, we measured trehalose levels as a percentage of dry cell mass before the first and after the last cycle (Figure 3H). The results indicated similar results among all media with a slightly smaller value for 2SMol. Before the first cycle, trehalose represented around 2.5% of the total dry mass, increasing to about 16% after the last. In contrast to the molasses investigated here, Raghavendran et al. [11] reported that trehalose content after six cycles was closer to the values observed for our 2SMol medium, representing about 11% of cellular dry mass. It is noteworthy that Raghavendran et al. [11] used sugarcane molasses obtained from *São Manoel* (São Manuel, Brazil) and *São Martinho* mill (Pradópolis, Brazil).

The comparison between the 2SMol medium and different real molasses performed here, considering all measurements and calculations made, indicate so far that the 2SMol formulation is capable of adequately reproducing sugarcane molasses, with a 100% defined composition. For specific situations or applications, it is possible to alter the original 2SMol composition, similar to what has been reported in the context of wine fermentations, in which particular characteristics of different grapes are explored by using different versions of the synthetic wine must medium reported by Bely et al. [12].

### Case study: How do nitrogen concentrations affect fed-batch fermentations in industrial-like conditions?

According to the literature, industrial alcoholic fermentations carried out in Brazilian sugar cane biorefineries lead to the highest reported ethanol yields per sugar consumed within all other substrates used [9, 41]. Several factors, such as the use of high cell concentrations and fed-batch operation with cell recycling, contribute to this outcome [4]. However, one of the hypotheses discussed by some authors is that the nitrogen content also plays a key role in this outcome since sugarcane feedstock are very poor in this nutrient, presumably being only sufficient to support 5 to 10% of biomass increase during each fermentation cycle [2, 4, 42]. In this scenario, the increase in cell biomass due to nitrogen availability is sufficient to replace losses of cell biomass caused by cell death and post-fermentation treatments (e.g., acid treatment, centrifugation, excess yeast cell biomass withdrawal).

We decided to take advantage of our fully defined synthetic molasses to investigate the effect of nitrogen availability on yeast performance in an industrially relevant setup. The 2SMol formulation was prepared with two different compositions in which the total nitrogen content was either zero (0SMol) or four times the amount (4SMol). The ratio between ammonia and amino acid nitrogen was kept constant. Thus, the total nitrogen concentrations were 0, 602.7, and 2401.6 mg.L^-1^ in 0SMol, 2SMol, and 4SMol, respectively.

Initially, results for CO_2_ loss normalized to wet cell mass were plotted as in previous research [11, 39]. Normalization ensures the biomass increase/decrease is considered during the evaluation of the results, and there is no over/underestimation of the outcomes [11]. Additionally, the CO_2_ loss could be directly related to sugar consumption along the fermentation cycle. The results of CO_2_ loss per gram of wet cell indicated a similar consumption rate among the three studied media (slope of the CO_2_ loss curve) in the first six hours of each cycle (Figure 4A). The final results obtained were different, in which a higher CO_2_ production per gram of cell was observed for the composition without nitrogen (15.0 ± 0.45 mmol.g^-1^) and decreased production for the 4SMol composition (12.27 ± 0.45 g.g^-1^), in comparison to the reference 2SMol composition (13.18 ± 0.23 g.g^-1^).

**Figure 4.**
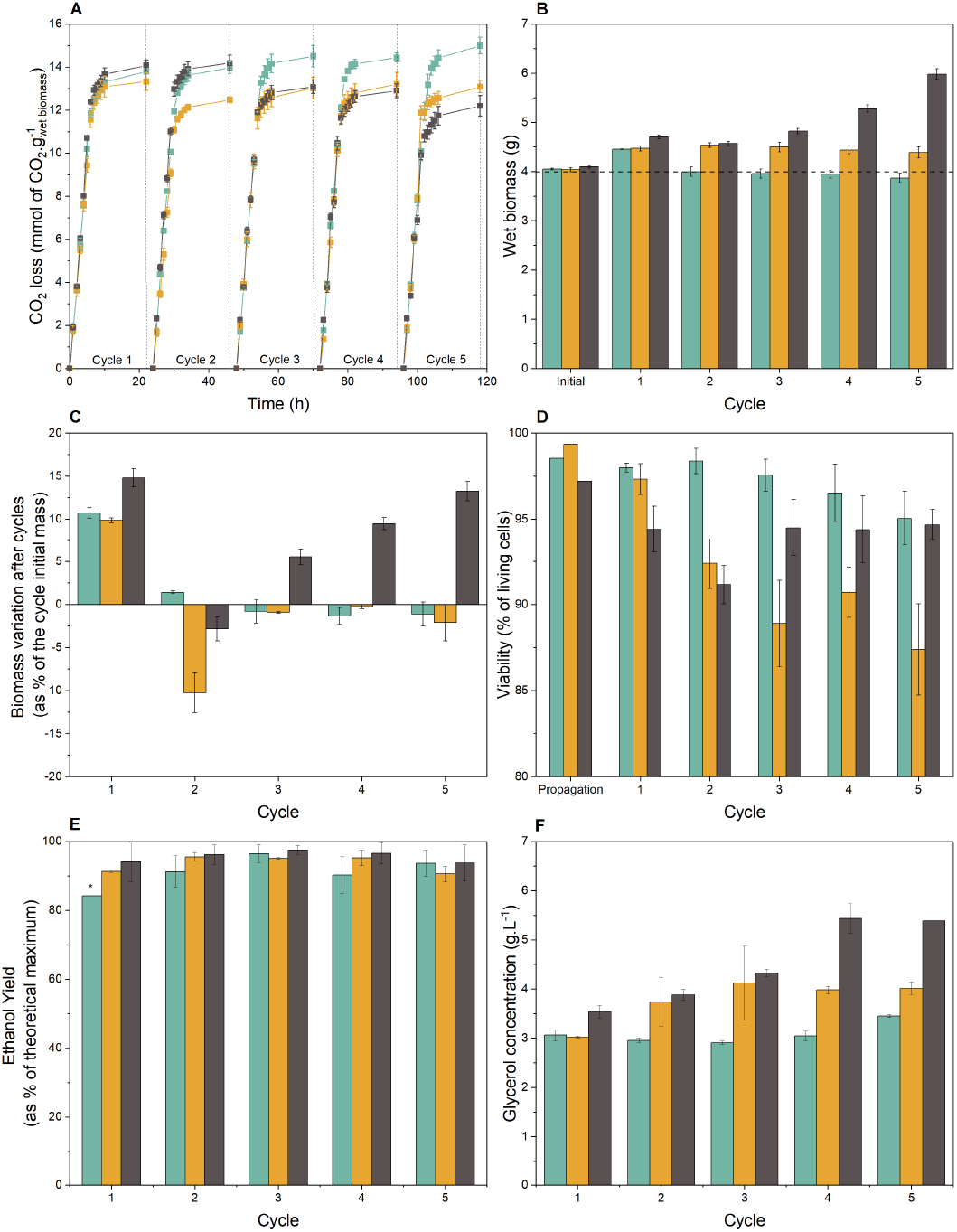
Nitrogen concentration effect in performance of *S. cerevisiae* PE-2 in the basal-defined synthetic molasses (2SMol), in the absence of nitrogen (0SMol), and 4-fold nitrogen concentrations in comparison with 2SMol (4SMol) in a scaled-down sugarcane biorefinery throughout five consecutive cycles. The fermentations were performed in triplicates. The initial yeast biomass was 4 g, acid treatment (pH set to 2.5, 1 h incubation at room temperature) was performed before each new cycle of fermentation. The tubes were incubated at 32 °C for 10 h and weighted hourly. The tubes were kept at room temperature (c.a. 20°C) overnight until the beginning of the next cycle. Orange (2SMol); Blue (0SMol); Grey (4SMol). (A) CO_2_ profiles (CO_2_ loss in mmol.g of wet biomass) over five fermentation cycles. (B) Overall biomass accumulation (g of wet biomass). (C) Biomass variation (%) represented as percentage of the cycle initial wet yeast mass. (D) Viability (as percentage of living cells, %) values by the methylene blue staining technique [44]. (E) Ethanol yield (as percentage of theoretical maximum 0.511 g of ethanol.g of TRS^-1^, %) calculated based on [11] and [35] (*) mean value for a duplicate. (F) Glycerol concentrations (g.L^-1^).

The values obtained for CO_2_ loss corroborated the data of biomass accumulated over the fermentation cycles (Figure 4B). The three media investigated started with 4 g of wet biomass with similar behaviors after the first cycle and distinct behaviors later on. After the first cycle, the resulting masses obtained were 4.45 ± 0.01, 4.47 ± 0.05, and 4.70 ± 0.04 g of wet cells, resulting in percentage variations close to 10 and 15 % of the initial fermentation mass (Figure 4C). Even in the absence of nitrogen, the initial increase can likely be explained by an intracellular accumulation of nutrients in the propagation stages preceding the fermentation cycles. In the 0SMol medium, cell biomass decreased in all the following cycles, leading to a total reduction of 4.6% of the initial mass. The viability for this medium also decreased from 99.5% to 87.4%, following the same downward trend of the total biomass (Figure 4D). The opposite pattern was observed for the medium with high nitrogen concentrations (4SMol). After an initial increase, the biomass was reduced at the end of the second cycle and significantly increased until the end of fermentation, finishing the fifth cycle with 5.98 ± 0.11 g of cells (45.97 ± 2.0%). Viability in 4SMol remained virtually stable over the cycles. As expected, glycerol titters were correlated to wet cell biomass, due to the necessity of reoxidazing excess NADH formed during biosynthesis [45]. Glycerol concentrations for 4SMol were higher cycle after cycle, when compared to 2SMol, reaching 5.4 g.L^-1^ at the end of the fifth fermentation cycle (Figure 4F). On the other hand, 0SMol presented the lowest glycerol contents with stabilization of production close to 3 g.L^-1^ and a slight increase in the last fermentation cycle.

Ethanol production was also compared and remained similar throughout fermentation, for the three media investigated. 0SMol led to low yield values, when compared to the theoretical maximum in the first fermentation cycle but reached values above 90% in the following cycles (Figure 4E). In general, the Brazilian industrial operation uses 2 to 3 fermentation cycles daily, which can reach 200 days of operation with cell recycling during each yearly production period [9]. From an industrial point of view, obtaining high ethanol yields with reduced cell viability and total biomass is not very advantageous, making the operation unfeasible for long periods and with the need for external replacement of cells. In this case, the media with nitrogen sources (2SMol and 4SMol) showed more balanced performances in the biomass/ethanol production ratio. Both 2SMol and 4SMol showed stabilization or growth of total biomass and remained with high ethanol yields in most cycles (c.a. >95%), indicating no change in production with increased nitrogen concentrations.

## CONCLUSIONS

This work illustrates the development of a fully defined synthetic molasses medium, that is based on a previously published semi-defined formulation. The proposed medium is conveniently prepared from some stock solutions and was validated in a scaled-down sugarcane biorefinery model, against real molasses-based media. In addition, this proposed formulation can be used to investigate the effects of relevant nutrients, as well as toxic compounds, on bioprocess performance. We thus hope the 2SMol formulation will be valuable to researchers both in academia and industry to obtain new insights and developments in industrial yeast biotechnology, and to provide reproducibility among different laboratories interested in sugarcane molasses-based processes.

## DECLARATIONS

### AVAILABILITY OF DATA AND MATERIALS

The datasets used and/or analysed during the current study are available from the corresponding author on reasonable request.

### AUTHORS’ CONTRIBUTIONS

**Kevy Eliodório:** Conceptualization, Methodology, Investigation, Validation, Formal analysis, Writing – Original Draft, Visualization. **Gabriel de Gois e Cunha:** Conceptualization, Methodology, Investigation, Writing - Review & Editing. **Felipe Lino:** Conceptualization, Writing - Review & Editing. **Morten Sommer:** Conceptualization, Writing - Review & Editing. **Andreas Gombert:** Conceptualization, Writing - Review & Editing. **Reinaldo Giudici:** Conceptualization, Resources, Writing - Review & Editing, Supervision, Funding acquisition. **Thiago Basso:** Conceptualization, Formal analysis, Resources, Writing - Review & Editing, Supervision, Project administration, Funding acquisition.

