## Supplementary files for "A Fully Defined Synthetic Medium Mimicking Sugar Cane Molasses"

* Corresponding author

Telephone: + 55 11 30912260

### Supplementary data

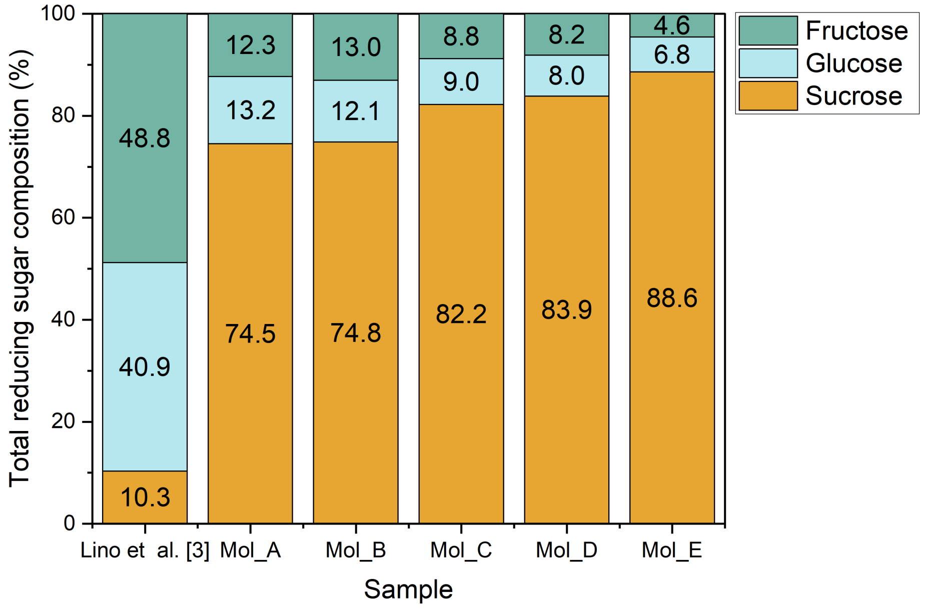


Figure S1 – Total reducing sugar content (%) of five sugarcane molasses samples (Mol_A, Mol_B, Mol_C, Mol_D, and Mol_E) and the semi-defined synthetic molasses proposed by Lino et al. [3]

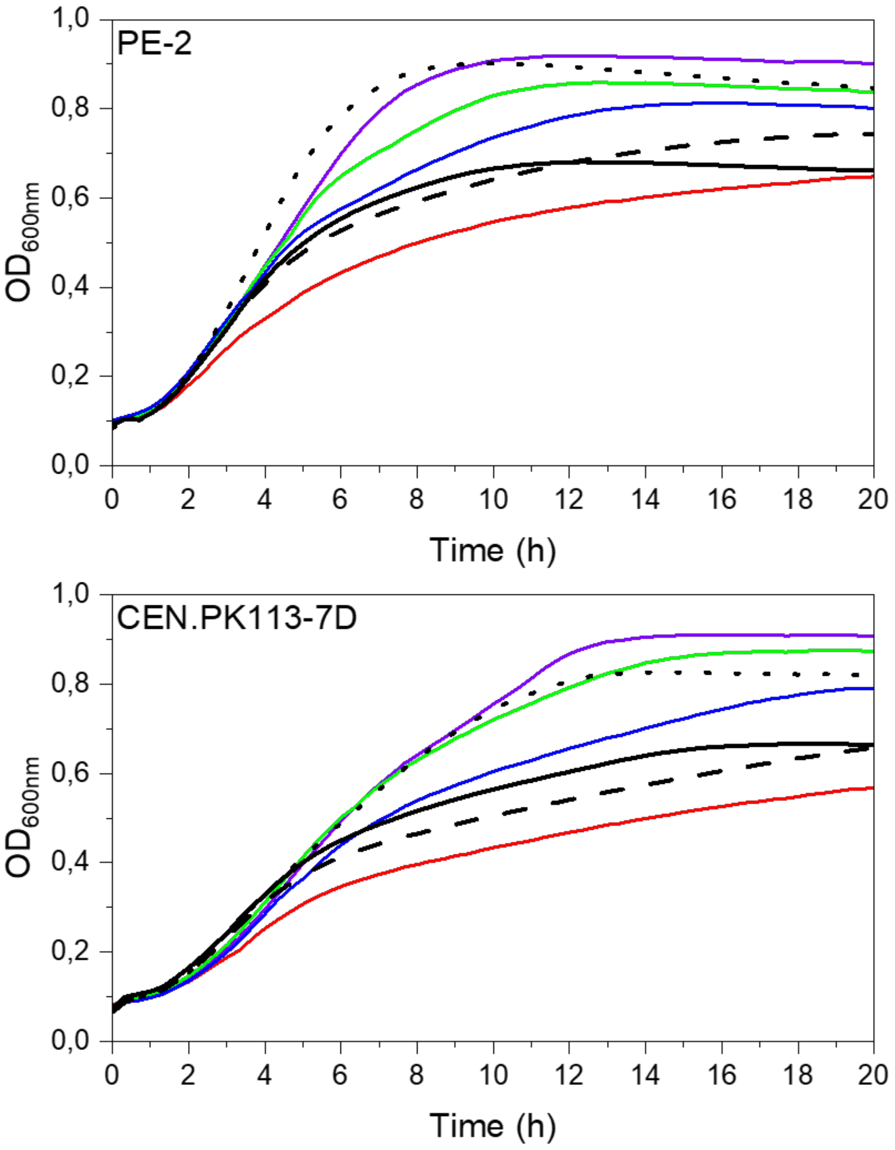


Figure S2 - Growth kinetics of *S. cerevisiae* strains in molasses (black lines, Mol_A: solid line, Mol_B: dotted line, and Mol_D: dashed line) and modified versions of 2SMol (0%: red, 25%: blue, 50%: green, and 100% amino acids: purple) media.

Table S1 - Compositional variations concerning 2SMolAA25 used in microplate assays to assess effect on yeast growth parameters with coded values in brackets.

| Condition | Concentration ratio of 2SMol | |
|---|---|---|
| | Lower | Upper |
| 2SMolAA25 | - | |
| Inorganic nitrogen | 0 (-1) | 2 (+1) |
| Organic Acids | 0 (-1) | 2 (+1) |
| Trace elements | 0.1 (-1) | 10 (+1) |
| Vitamins | 0.1 (-1) | 2 (+1) |
| Mg & K | 0.2 (-1) | 2 (+1) |
| Calcium | 0 (-1) | 2 (+1) |
| Organic nitrogen | 0 (-1) | 0.5 (+1) |
| Phosphate | 0.5 (-1) | 2 (+1) |

*The tested conditions are presented as the ratio between theses variations for each nutritional group and the composition of 2SMol containing 25% of amino acids (2SMolAA25)

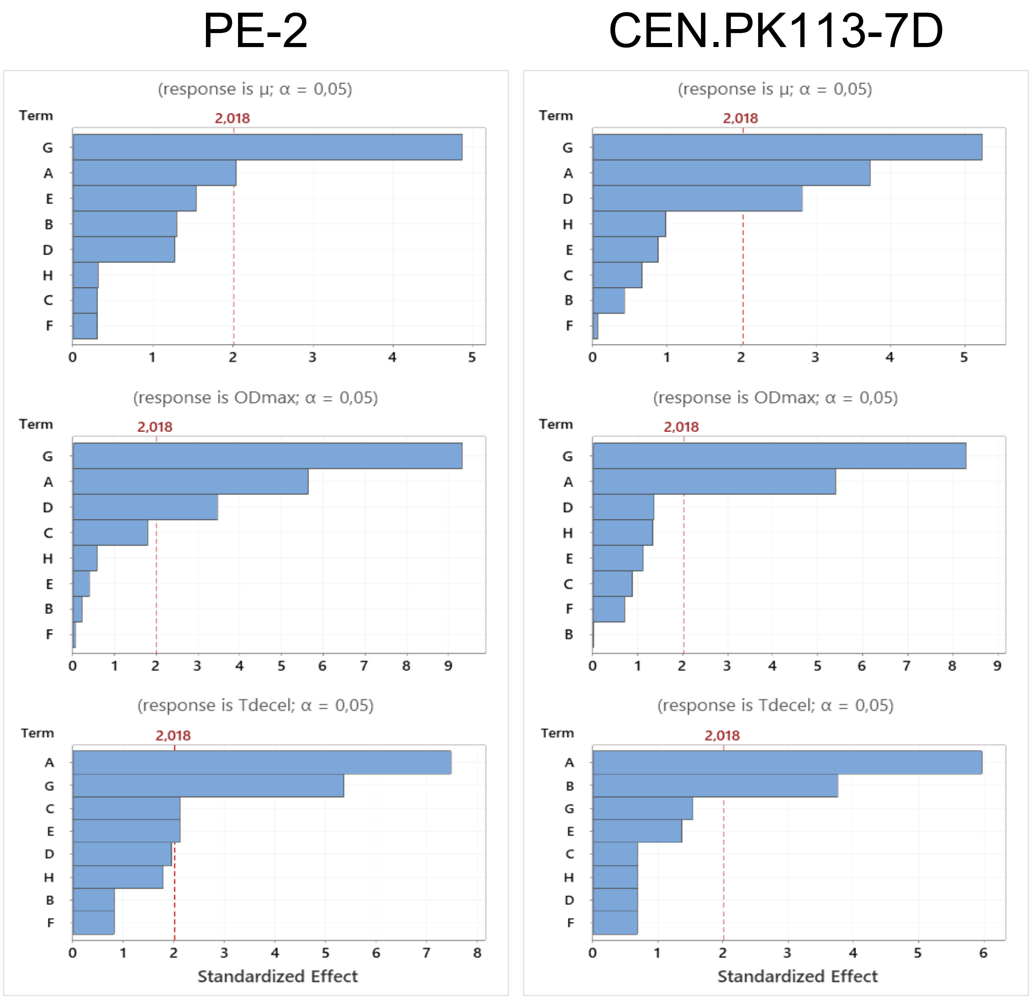


Figure S3 - Pareto chart of the standardized effects for CEN.PK113-7D and PE-2 in 2SMolAA25 with variations in the concentration of nutritional groups. (A) Inorganic nitrogen, (B) Organic acids, (C) Trace elements, (D) Vitamins, (E) Mg and K, (F) Calcium, (G) Organic nitrogen, and (H) Phosphate.

Table S2 - Growth parameters for *S. cerevisiae* strains PE-2 and CEN.PK113-7D using lower and higher levels of eight nutritional groups of 2SmolAA25.

| **PE-2** | | | | | | |
|---|---|---|---|---|---|---|
| Parameter | μ ($h^{-1}$) | | $OD_{Max}$ | | $T_{decel}$ (h) | |
| Level | Lower | Higher | Lower | Higher | Lower | Higher |
| **2SMolAA25** | 0.416 | | 0.823 | | 12.22 | |
| **Inorganic nitrogen** | 0.428 | 0.4 | 0.812 | 0.688 | 8.22 | 13.33 |
| **Organic Acids** | 0.432 | 0.414 | 0.796 | 0.8 | 10.67 | 10.11 |
| **Trace elements** | 0.407 | 0.403 | 0.778 | 0.817 | 11.78 | 10.34 |
| **Vitamins** | 0.393 | 0.411 | 0.736 | 0.812 | 13 | 11.67 |
| **Mg & K** | 0.433 | 0.412 | 0.789 | 0.798 | 12 | 10.56 |
| **Calcium** | 0.425 | 0.421 | 0.81 | 0.809 | 12.22 | 11.67 |
| **Organic nitrogen** | 0.357 | 0.424 | 0.682 | 0.889 | 13.33 | 9.67 |
| **Phosphate** | 0.419 | 0.415 | 0.787 | 0.8 | 11.56 | 12.78 |
| Mol_B | 0.496 | | 0.902 | | 5.78 | |
| Mol-A | 0.426 | | 0.682 | | 8.56 | |
| Mol_D | 0.417 | | 0.746 | | 17.56 | |
| **CEN.PK113-7D** | | | | | | |
| Parameter | μ ($h^{-1}$) | | $OD_{Max}$ | | $T_{decel}$ (h) | |
| Level | Lower | Higher | Lower | Higher | Lower | Higher |
| **2SMolAA25** | 0.387 | | 0.833 | | 17 | |
| **Inorganic nitrogen** | 0.404 | 0.367 | 0.857 | 0.689 | 13.11 | 17 |
| **Organic Acids** | 0.382 | 0.387 | 0.813 | 0.812 | 14.22 | 16.67 |
| **Trace elements** | 0.367 | 0.373 | 0.852 | 0.825 | 16.11 | 16.56 |
| **Vitamins** | 0.348 | 0.377 | 0.836 | 0.878 | 16.89 | 16.45 |
| **Mg & K** | 0.388 | 0.379 | 0.84 | 0.806 | 16.89 | 16 |
| **Calcium** | 0.384 | 0.385 | 0.818 | 0.84 | 16.89 | 16.45 |
| **Organic nitrogen** | 0.335 | 0.388 | 0.638 | 0.896 | 17 | 16 |
| **Phosphate** | 0.365 | 0.375 | 0.797 | 0.838 | 16.22 | 16.67 |
| Mol_B | 0.390 | | 0.827 | | 10.78 | |
| Mol_A | 0.380 | | 0.666 | | 14.56 | |
| Mol_D | 0.370 | | 0.716 | | 26 | |

* Standard deviations are not shown, given that they represented less than 8% of each mean value.

Table S3 - Investigated factors for the $3^3$ factorial design and their levels with coded values in brackets.

| Investigated factors | Factor levels in % of 2SMol content | | |
|---|---|---|---|
| Organic nitrogen | 0 (-1) | 37,5 (0) | 75 (+1) |
| Inorganic nitrogen | 0 (-1) | 100 (0) | 200 (+1) |
| Vitamin | 0 (-1) | 100 (0) | 200 (+1) |

Table S4 - Model coefficients for maximum specific growth rate (μ) obtained from the $3^3$ factorial design for yeast strains PE-2 and CEN.PK113-7D. (1) Organic nitrogen, (2) Inorganic nitrogen, (3) Vitamins.

| Term | PE-2 | | | | CEN.PK113-7D | | | |
|---|---|---|---|---|---|---|---|---|
| | Coefficient | Standard Error | t-Value | p-Value | Coefficient | Standard Error | t-Value | p-Value |
| $b_0$ | 0.424 | 0.006 | 68.820 | 0.000 | 0.401 | 0.008 | 53.100 | 0.000 |
| $b_1$ | 0.044 | 0.003 | 15.540 | 0.000 | 0.046 | 0.004 | 13.030 | 0.000 |
| $b_2$ | 0.026 | 0.003 | 9.130 | 0.000 | 0.032 | 0.004 | 9.130 | 0.000 |
| $b_3$ | 0.023 | 0.003 | 8.230 | 0.000 | 0.035 | 0.004 | 9.890 | 0.000 |
| $b_{11}$ | -0.037 | 0.005 | -7.550 | 0.000 | -0.048 | 0.006 | -7.860 | 0.000 |
| $b_{22}$ | -0.023 | 0.005 | -4.680 | 0.000 | -0.033 | 0.006 | -5.460 | 0.000 |
| $b_{33}$ | -0.024 | 0.005 | -4.810 | 0.000 | -0.031 | 0.006 | -5.170 | 0.000 |
| $b_{12}$ | -0.030 | 0.003 | -8.510 | 0.000 | -0.048 | 0.004 | -11.140 | 0.000 |
| $b_{13}$ | 0.006 | 0.003 | 1.730 | 0.089 | 0.008 | 0.004 | 1.870 | 0.065 |
| $b_{23}$ | 0.011 | 0.003 | 3.010 | 0.004 | 0.006 | 0.004 | 1.330 | 0.188 |
| $R^2$ | | 89.08% | | | | 89.40% | | |
| $R^2$ adjusted | | 87.69% | | | | 88.05% | | |
| $R^2$ predicted | | 85.57% | | | | 86.34% | | |

Table S5 - Model coefficients for maximum absorbance ($OD_{max}$) obtained from the $3^3$ factorial design for yeast strains PE-2 and CEN.PK113-7D. (1) Organic nitrogen, (2) Inorganic nitrogen, (3) Vitamins.

| Term | PE-2 | | | | CEN.PK113-7D | | | |
|---|---|---|---|---|---|---|---|---|
| | Coefficient | Standard Error | t-Value | p-Value | Coefficient | Standard Error | t-Value | p-Value |
| $b_0$ | 0.914 | 0.017 | 55.310 | 0.000 | 0.903 | 0.021 | 44.040 | 0.000 |
| $b_1$ | 0.159 | 0.008 | 20.840 | 0.000 | 0.167 | 0.009 | 17.580 | 0.000 |
| $b_2$ | 0.076 | 0.008 | 9.890 | 0.000 | 0.074 | 0.009 | 7.840 | 0.000 |
| $b_3$ | 0.032 | 0.008 | 4.230 | 0.000 | 0.044 | 0.009 | 4.650 | 0.000 |
| $b_{11}$ | -0.081 | 0.013 | -6.100 | 0.000 | -0.056 | 0.016 | -3.400 | 0.001 |
| $b_{22}$ | -0.042 | 0.013 | -3.160 | 0.002 | -0.055 | 0.016 | -3.370 | 0.001 |
| $b_{33}$ | -0.017 | 0.013 | -1.280 | 0.205 | -0.006 | 0.016 | -0.360 | 0.722 |
| $b_{12}$ | -0.066 | 0.009 | -7.060 | 0.000 | -0.103 | 0.012 | -8.840 | 0.000 |
| $b_{13}$ | 0.034 | 0.009 | 3.640 | 0.001 | 0.029 | 0.012 | 2.480 | 0.016 |
| $b_{23}$ | 0.014 | 0.009 | 1.450 | 0.150 | 0.016 | 0.012 | 1.370 | 0.176 |
| $R^2$ | 90.34% | | | | 82.35% | | | |
| $R^2$ adjusted | 89.11% | | | | 82.37% | | | |
| $R^2$ predicted | 87.49% | | | | 79.69% | | | |

Table S6 - Model coefficients for deceleration time ($t_{decel}$) obtained from the $3^3$ factorial design for yeast strains PE-2 and CEN.PK113-7D. (1) Organic nitrogen, (2) Inorganic nitrogen, (3) Vitamins.

| Term | PE-2 | | | | CEN.PK113-7D | | | |
|---|---|---|---|---|---|---|---|---|
| | Coefficient | Standard Error | t-Value | p-Value | Coefficient | Standard Error | t-Value | p-Value |
| $b_0$ | 11.137 | 0.378 | 29.480 | 0.000 | 18.274 | 0.949 | 19.260 | 0.000 |
| $b_1$ | -4.315 | 0.175 | -24.670 | 0.000 | -4.889 | 0.439 | -11.130 | 0.000 |
| $b_2$ | -2.297 | 0.175 | -13.130 | 0.000 | -3.704 | 0.439 | -8.430 | 0.000 |
| $b_3$ | -0.766 | 0.175 | -4.380 | 0.000 | -4.463 | 0.439 | -10.160 | 0.000 |
| $b_{11}$ | 0.352 | 0.303 | 1.160 | 0.249 | 1.148 | 0.761 | 1.510 | 0.136 |
| $b_{22}$ | 0.037 | 0.303 | 0.120 | 0.903 | -0.444 | 0.761 | -0.580 | 0.561 |
| $b_{33}$ | 0.704 | 0.303 | 2.320 | 0.023 | 2.871 | 0.761 | 3.770 | 0.000 |
| $b_{12}$ | 0.111 | 0.214 | 0.520 | 0.606 | 1.482 | 0.538 | 2.750 | 0.007 |
| $b_{13}$ | -0.620 | 0.214 | -2.900 | 0.005 | -4.084 | 0.538 | -7.590 | 0.000 |
| $b_{23}$ | -0.796 | 0.214 | -3.720 | 0.000 | -0.806 | 0.538 | -1.500 | 0.139 |
| $R^2$ | | 92.13% | | | | 87.59% | | |
| $R^2$ adjusted | | 91.12% | | | | 86.02% | | |
| $R^2$ predicted | | 89.60% | | | | 84.04% | | |

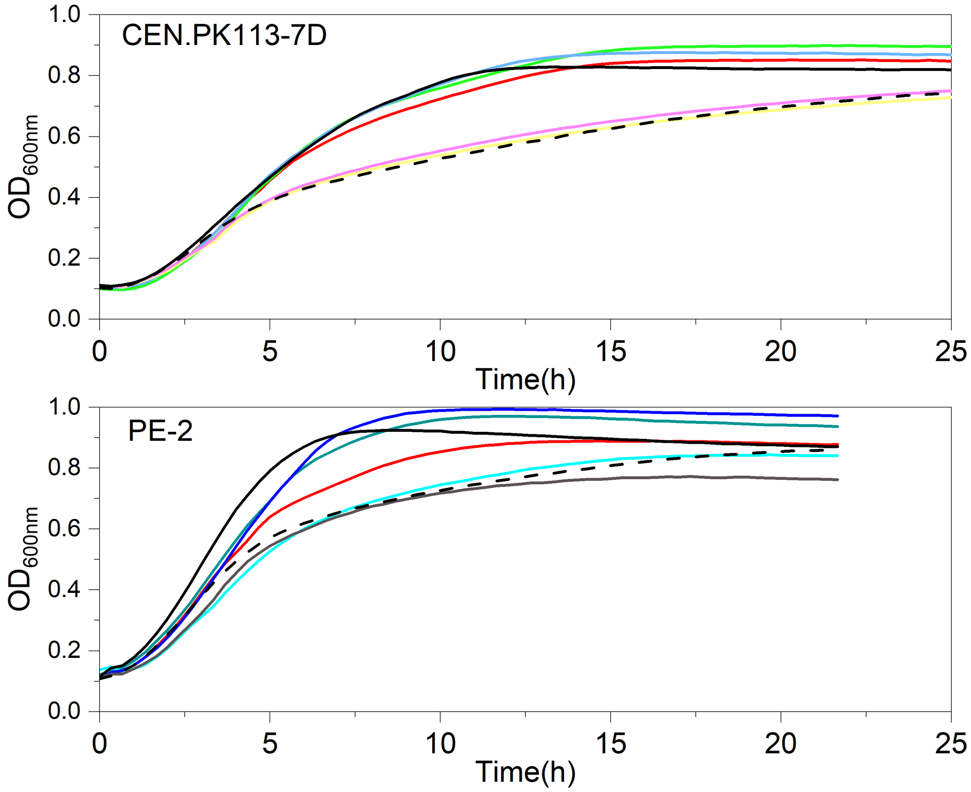


Figure S4 - Growth kinetics of *S. cerevisiae* strains CEN.PK 113-7D and PE-2 in: molasses (black lines, Mol_B solid, and Mol_D dashed) and modified versions of 2SMol [% of Organic nitrogen, Inorganic Nitrogen, vitamins in 2SMol]: [37.5, 100, 100] red line, [37.5, 200, 200] dark green, [37.5, 0, 200] light blue, [75, 100, 100] marine blue, [0, 200, 100] grey, [37.5, 100, 200] green, [37.5, 200, 100] light blue, [0, 100, 100] yellow, [0, 100, 200] pink
